# Causally investigating cortical dynamics and signal processing by targeting natural system attractors with precisely timed stimulation

**DOI:** 10.1101/454538

**Authors:** Dmitriy Lisitsyn, Udo A. Ernst

## Abstract

Electrical stimulation is a promising tool for interacting with neuronal dynamics to identify neural mechanisms that underlie cognitive function. Since effects of a single short stimulation pulse typically vary greatly and depend on the current network state, many experimental paradigms have rather resorted to continuous or periodic stimulation in order to establish and maintain a desired effect. However, such an approach explicitly leads to forced and ‘unnatural’ brain activity. Further, continuous stimulation can make it hard to parse the recorded activity and separate neural signal from stimulation artifacts. In this study we propose an alternate strategy: by monitoring a system in realtime, we use the existing preferred states or attractors of the network and to apply short and precise pulses in order to switch between its preferred states. When pushed into one of its attractors, one can use the natural tendency of the system to remain in such a state to prolong the effect of a stimulation pulse, opening a larger window of opportunity to observe the consequences on cognitive processing. To elaborate on this idea, we consider flexible information routing in the visual cortex as a prototypical example. When processing a stimulus, neural populations in the visual cortex have been found to engage in synchronized gamma activity. In this context, selective signal routing is achieved by changing the relative phase between oscillatory activity in sending and receiving populations (communication through coherence, CTC). In order to explore how perturbations interact with CTC, we investigate a biophysically realistic network exhibiting similar synchronization and signal routing phenomena. We develop a closed-loop stimulation paradigm based on the phase-response characteristics of the network and demonstrate its ability to establish desired synchronization states. By measuring information content throughout the model, we evaluate the effect of signal contamination caused by the stimulation in relation to the magnitude of the injected pulses and intrinsic noise in the system. Finally, we demonstrate that, up to a critical noise level, precisely timed perturbations can be used to artificially induce the effect of attention by selectively routing visual signals to higher cortical areas.

## 2 Introduction

With evolving technology, new and promising techniques to interfere with the brains natural activity have played a crucial role in moving from correlational to causal links between neuronal activity and behavior [46, 30, 18]. Crucially, the same techniques are used clinically to treat pathological injuries and disorders [32, 6, 19, 7]. The development of perturbation technology, among many others, includes ablations of cortical and subcortical targets, chemical lesions, reversible inactivations, transcranial direction current stimulation (tDCS), transcranial magnetic stimulation (TMS), intracortical microstimulation (ICMS), and finally the fairly recent and exciting optogenetic techniques [57]. This advancement of tools has provided increasingly higher temporal and spatial perturbation precision, allowing for more intricate control over neural activity, which in turn has supported progressively stronger conclusions about the neuronal mechanisms underlying cognition.

While non-invasive techniques such as tDCS and TMS ease clinical applicability, the effects of their stimulation unfortunately lack spatial precision. Invasive techniques, in particular, ICMS and optogenetics allow for precise temporal and spatial resolution, providing the ability to deliver a single short and temporally precise perturbation at a precise location in the brain, which in turn, should greatly increase the ability to accurately affect and control neural circuits. However, the effect of such a single short perturbation can be very short-lived and, crucially, it can vary greatly in dependence on the state of the neural system at the pulse onset. Because of this, many perturbation paradigms have opted to either use a very strong pulse, essentially resetting and disrupting the activity of the target network, or to use a continuous or repetitive-pulse stimulation in order to establish and maintain a desired effect. For instance, a series of seminal studies [15, 40] entrained a local population in the barrel cortex of mice with a rhythmic optogenetic train of pulses at 40 Hz. By delivering a vibrissa stimulation at different phases of the entrained populations cyclic activity, the researchers showed that the neural populations response as well as the rodents behavioral performance depends on the phase at which the whisker stimulation stimulus arrives to the population. In a more recent study, Ni et al 2016 used a similar technique to show how an optogenetically induced neural rhythm modulates the gain of spike responses and behavioral reaction times in response to visual stimuli in cats [36].

Using continuous stimulation serves its role as a powerful research tool, however it also brings up a number of concerns. First, in some cases, stimulation can effectively destroy and suppress any ongoing local processing [30]. Even if it does not lead to full suppression, in addition to achieving a desired effect, continuous stimulation may interfere and contaminate the relevant neural signals. Further, in many cases, when analyzing the activity recorded during the stimulation, it becomes hard, if not unfeasible, to separate the stimulation artifacts from the relevant neural data. Finally, such an approach explicitly forces the neural system to remain in some desired network state, resulting in artificial dynamics and making it questionable what we learn about processing during natural activity.

In this study, we propose to use an alternate strategy. Rather than using continuous stimulation in order to sustain a desired state of the neural network, we wish to utilize a single precise pulse in order to push the system into one of its (potentially) existing preferred states [48]. If the network is pushed into one of its attractors, the natural tendency of the system to remain in such a state extends the duration of the effect of the pulse, which opens up a larger window of opportunity to observe the consequences on cognitive processing. Crucially, it becomes necessary to monitor the system in real time in order to be aware of the systems state and to deliver just the right stimulation at just the right time, resulting in a closed-loop paradigm.

The brains rhythmic activity and synchronization phenomena provide a perfect test-bed for our approach. Brain rhythms have been at the center of neuroscience research since they were first observed with the invention of EEG (electroencephalography) over a century ago [17]. Neural oscillations are generated at specific frequencies, coexisting with background noise (non-oscillatory) activity. They can be observed at multiple scales, from the activity of a single neuron to the coordinated output of large neuronal networks [50]. Further, distinct neural populations can entrain each other, exhibiting coupled states and synchronized activity [41, 50]. Research has shown that such neural synchrony plays a crucial role that underlies many cognitive processes, such as perceptual grouping [38], working memory [37] and information routing of signals throughout the brain [21]. Clinically, abnormal levels of synchrony have been linked to pathological disorders, such as schizophrenia, autism, Alzheimers and Parkinsons [49].

Approaching the brains rhythmic activity and synchronization phenomena from a perspective of non-linear dynamics provides useful inferences on neural oscillator activity [26]. First, oscillatory synchronization collapses the normally high dimensional dynamics of neural dynamics into a low dimensional set of attractor states. Further, if a neural system can be modeled using self-sustained, weakly chaotic oscillators, a perturbation inserted at a specific phase of a cycle would evoke a consistent phase-shift in the oscillators activity an effect that is captured by a phase-response-curve (PRC) [39, 13]. Numerous experimental studies have found evidence for PRCs in vitro [2] and as well as in vivo [52, 51, 54].

In the present study, through a modeling approach, we develop a method to explore the feasibility of utilizing PRCs in order to shift the synchronization of a system into a desired state. First, we choose to model selective information routing in the visual cortex, between V1 and V4 cortical areas. A prominent mechanism explaining how information routing occurs, communication through coherence (CTC), relies on the inherent oscillatory dynamics of neural activity and postulates that neural populations establish favorable and unfavorable information routing states through frequency-specific phase-locking [21, 22]. In support of this hypothesis, experimental studies have shown strong evidence for gamma-band synchronization between sending V1 and receiving V4 neural populations during a visual attention task [10, 24]. Once a favorable synchronization state is established, rhythmic bursts of V1 spikes arrive to V4 during its excitability peaks, increasing the likelihood that further spikes are evoked leading to effective signal routing. On the other hand, if the V1 and V4 populations establish an unfavorable phase state relationship, the V1 spikes arrive to V4 during the excitability troughs and hence should fail or at least be less effective in evoking further activity.

We begin with a model of an isolated neural oscillator and then expand to biophysically realistic system of multiple coupled populations, constructed to exhibit the synchronization and information routing phenomena observed in the visual cortex. We explicitly measure the information content in the model to evaluate the effect of signal contamination caused by the stimulation in relation to the magnitude of the injected pulses and intrinsic noise level of the system. Further, we vary the background noise level to investigate how increased stochasticity affects the phase-response properties the system and hence our ability to control it. We demonstrate that up to a critical noise level, precisely timed perturbations can be used to ‘simulate’ the effect of attention by selectively routing a visual signal to higher cortical areas and identify optimal pulse strengths required to achieve this goal.

## 3 Results

In the first part of this section, we present the model of a single cortical column as the basic building block of our framework. Further, we introduce the techniques needed to monitor oscillatory dynamics, and demonstrate how to use them to control single oscillators to maintain a desired system state. Taken together, these considerations pave the way for interacting with a more realistic, hierarchical cortical network in Part II of this section.

### 3.1 Part I: Stimulating a cortical column – basic concepts

#### 3.1.1 Cortical column model

##### Model structure and dynamics

For representing one cortical column, we construct a recurrent network with 800 excitatory and 200 inhibitory, conduction-based quadratic integrate-and-fire neurons. Their membrane potentials *V* evolve according to the differential equation

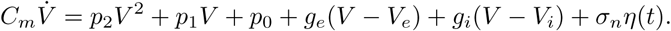

Here, *C*_*m*_ is the membrane capacitance, *V*_*e*_ and *V*_*i*_ are the reversal potentials and *g*_*e*_ and *g*_*i*_ the corresponding conductances for excitatory and inhibitory input currents, and *η*(*t*) represents a white noise term with magnitude *σ*_*n*_. If the membrane potential *V* crosses the threshold *V*_*thresh*_, a spike is generated and delivered to all connected neurons, and *V* is reset to *V*_*rest*_.

Synaptic term equations and all the relevant parameter values are presented in the Methods section. Connections exist from the inhibitory population to itself, with projection probability 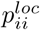 and corresponding delay 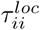, and from the inhibitory to the excitatory populations, with projection probability 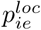 and corresponding delay 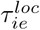 (Fig. 1A). Neuron and coupling parameters are set to emulate realistic neurons, in accordance with [4] (for details on the implementation and parameters see Methods section). In particular, delays are set to 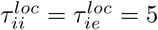 ms which allows the network to generate gamma frequency oscillations (Fig. 1B, upper panel) by means of an ING-mechanism [47].

**Figure 1.**
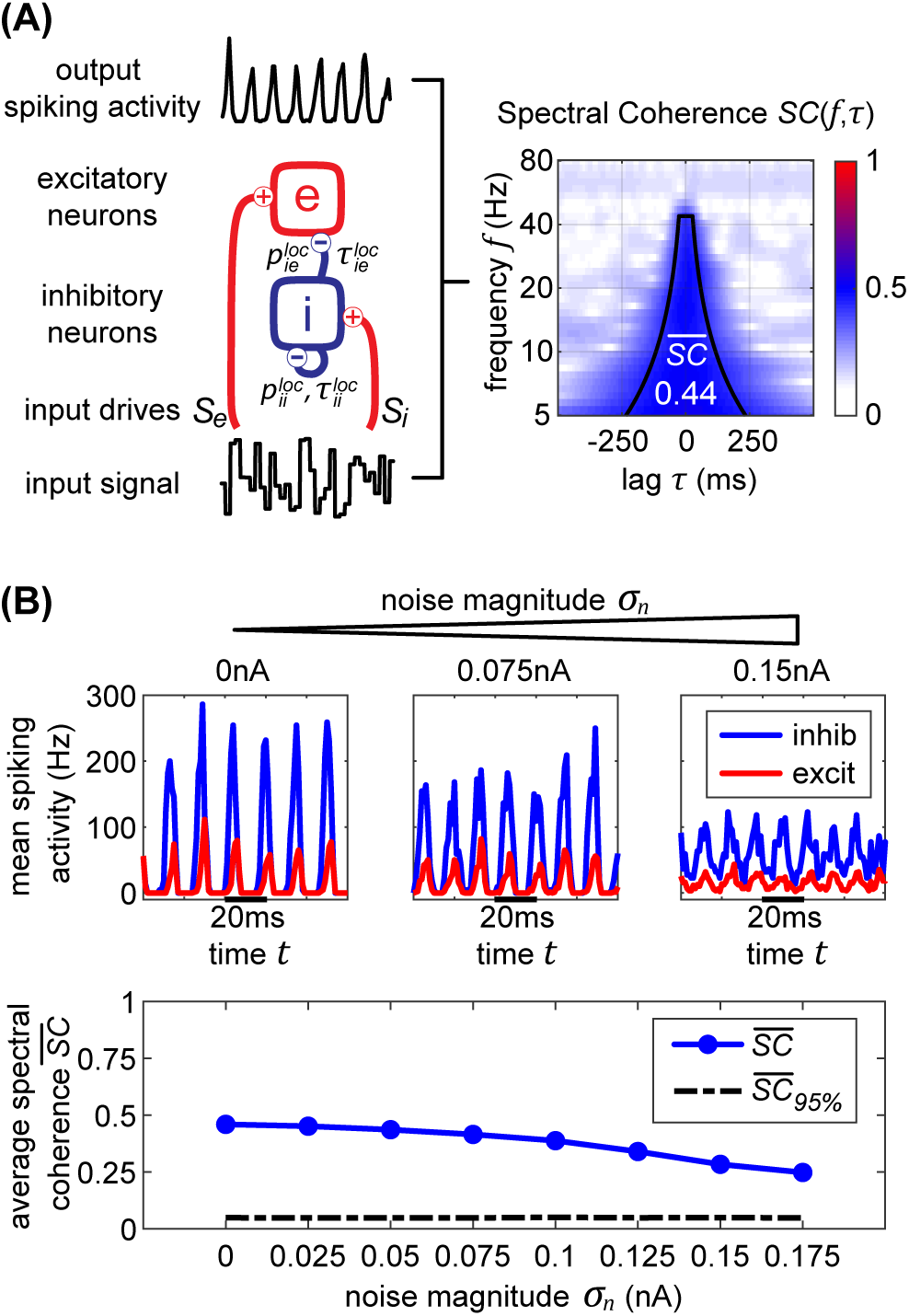
**Single oscillator model and activity.** **A.** A single cortical column is represented by a neural oscillator, consisting of an excitatory and an inhibitory neuron population. It is driven by an input signal containing a time-varying amplitude modulation. The inhibitory population projects onto itself and onto the excitatory neurons, resulting in an ING mechanism which produces cyclic population activity in the gamma frequency range (60-75Hz). To evaluate the signal routing ability of the network, we assess the spectral coherence SC(*f, τ*) between the input signal modulation and the excitatory output activity for different frequencies *f* and signal time lags *τ*. An cumulative input signal information measure 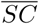 (white text) is computed by pooling across the relevant time-frequency range within the cone of interest (solid black lines). **B.** In the top row, we show samples of excitatory (red) and inhibitory (blue) population activity for increasing internal noise levels. At zero noise, oscillatory spiking activity is very regular - large population bursts are followed by periods of silence. With increasing noise, activity gets more irregular and less phase specific. In the bottom row, we show how input signal modulation contribution to the neural activity 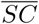 decreases with increasing internal noise *σ*_*n*_.

Both populations are driven by afferent connections delivering excitatory input with time-varying rates *S*_*e*_(*t*) and *S*_*i*_(*t*), realized by inhomogeneous Poisson processes. We scaled the mean rate and driving magnitude of the afferent input such that we achieve an average firing rate of 15 Hz for the excitatory units and 60 Hz for the inhibitory units, reflecting typical neuronal firing rates found in the cortex [53].

##### Quantifying stimulus representation

When interacting with a cortical network by external electric stimulation, we pursue two goals: Assessing the implied changes in dynamical network states, and quantifying the impact on function, i.e. the representation and processing of visual information. For the latter goal, we adopt a method which was used successfully to quantify selective signal transmission (’gating’) in dependence on the attentional state [27, 25]. The main idea of this method is to modulate the visual (input) signal by a random change in its amplitude (’flicker’), and to compare the output of a neural population with the input flicker signal by computing a frequency-resolved correlation termed spectral coherence (SC).

In our case, we modulated the external drive with mean rate 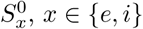

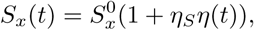

where *η*(*t*) was sampled from a uniform distribution between [−1, 1] with a rate of 1/*T*_*f*_. Emulating the signals used in [25] where a visual stimulus was presented at 100 frames per second, we set *T*_*f*_ = 10 ms.

The strength of drive modulation was set to *η*_*S*_ = 0.10. This modulation is passed onto the spiking rates of the driven neural populations (Fig. 1A bottom).

To evaluate the input modulation contribution to the neural activity, we utilize spectral coherence (SC). First, we compute the spectrograms of the input signal and the spike output using a wavelet transform with Morlet kernels. The transform yields complex valued coefficients *W*_*z*_(*f, t*) representing the amplitude and phase of a signal *z*(*t*) around the frequency band *f* at time *t*. By evaluating the normalized cross-correlation between the spectrograms of *x*(*t*) and *y*(*t*) we obtain the spectral coherence measure *C*_*xy*_(*f, τ*), where *f* is the frequency and *τ* is the delay between the two signals:

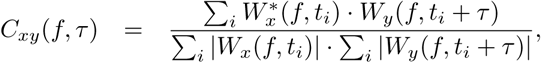

Due to the normalization terms in the denominator, the values of *C*_*xy*_ lie between zero and one.

If neurons are driven well by the external stimulus, experimental data [25] and model simulations (Fig.1A) reveal that the input signal can be tracked in the population activity of a cortical column in V1 (or V4) up to frequencies of about 45 Hz (or 25 Hz). Hence, in order to obtain a cumulative measure of input signal contribution to the neural activity, we defined a cone-of-interest whose upper frequency limit was selected to be at 45 Hz, and whose temporal range was defined as *±*7/6*f* around *τ* = *τ*_*xy*_, where *τ*_*xy*_ is the delay between input signal *x* and neural output *y*. We pooled across the relevant frequencies *f* and time lags *τ* within the cone of interest, to compute a single spectral coherence score 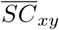.

##### Gamma oscillations and noise

In a typical experimental situation, it is impossible to assess the output signal of a specific neural population directly. Instead, the measurement is confounded by both, measurement noise and noise induced by background activity or by contributions from neighboring circuits. For interacting with the brain, it is therefore essential to quantify the impact of noise on the assessment of the current system state and to determine limits up to which successful control is still possible. We therefore introduced internal noise via the additional term *σ*_*n*_*η*(*t*) in equation 3.1.1 [20]. *η*(*t*) is 1/*f* noise with standard deviation equal to 1, making *σ*_*n*_ represent the magnitude of the noise. Every single neuron unit receives its own unique noise input. By changing the magnitude *σ*_*n*_, we control the overall level of noise in the entire system.

In Fig. 1B in the top three plots, we display model activity at different noise levels. With zero noise level we clearly see oscillations within the Gamma frequency range, with low jitter and high regularity and phase specificity – inhibitory and excitatory populations of neurons both evoke concentrated bursts of spikes followed by periods of silence. Increasing the noise renders oscillations more irregular and less phase-specific, and decreases peak amplitudes. Also, oscillation frequency increases from 60 Hz for the zero noise condition to 75 Hz for 0.15 nA. In order to maintain a stable cyclic activity with a constant frequency, the ratio of inhibitory and excitatory post-synaptic currents needs to stay consistent within each population of neurons [12]. Increasing the magnitude of noise inherently raises the firing rate of neurons. Since our units are recurrently coupled, a change in average firing rate upsets the inhibition-excitation ratio of the system, which results in a dramatic change in activity. To counteract this effect, for each noise level, we update the magnitude of driving rates *S*_*e*_ and *S*_*i*_ to provide just the right amount of input drive to excitatory and inhibitory units to maintain firing rates consistent with physiological evidence, i.e. an average of 15 Hz for excitatory and 60 Hz for inhibitory units (parameters see Methods section).

For noise levels of about 0.1 nA, we observe signals similar to physiological findings [25]. To cover a realistic range, we investigated noise levels from *σ*_*n*_ = 0 nA up to *σ*_*n*_ = 0.175 nA. Crucially, as can be expected, the input signal representation as quantified by 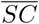 becomes worse with increasing noise, although it stays well above the significance level (Fig. 1B, bottom plot).

#### 3.1.2 Tracking oscillations and stimulation effects

##### Real-time phase tracking

For a targeted interaction with a neural system, we have to assess its internal state in real-time. In our case, the internal state is characterized by the current phase of an ongoing oscillation (in the Gamma frequency range). Consequently, we will have to determine this phase as precisely and timely as possible.

Tracking the phase of a signal in real-time imposes the constraint that only data from the past can be used for phase measurement, whereas the typical offline phase measurement algorithms rely on utilizing past and future data for an accurate estimate of the instantaneous phase at that time point. In our case, we use a phase-extraction scheme motivated by [16] that relies on an autoregression (AR) linear model for discrete time series prediction. The AR model has been found to perform well in forecasting noisy signals with power spectrum limited to certain frequencies [8], which makes it perfect for our data.

Before the AR model can be used, it must first be trained on data where no electrical stimulation was applied. We also use the same data to also determine the characteristics of gamma oscillations by calculating the power spectrum of the signal via a Morlet wavelet transform, from which we determine the location and halfway points of the gamma peak.

We then use the following three steps to determine the phase of a signal at time *t*: First, the forecasted signal is calculated. Then, we apply a zero-phase bandpass filter with bandstops at the halfway points found in the power spectrum for obtaining the gamma component of the signal without distorting its phase. Finally, the signal is passed through a Hilbert transform [9], providing us with the complex analytical signal. The argument of the analytical signal reveals the instantaneous gamma phase. The narrow range of the bandpass filter is necessary, since the instantaneous phase only becomes accurate and meaningful if the filter bandwidth is sufficiently narrow [35].

Crucially, this method also proves useful for offline data analysis. The same procedure from above can be applied to neural signals just prior to an input pulse to determine the phase of the ongoing oscillation before it is affected by any phase shifts or especially, recording artifacts. A similar method relying on AR was utilized specifically for this reason in [36].

##### Phase-response curves

Using stimulation pulses, our goal is ‘push’ a neural system towards particular states and quantify the impact of such a ‘configuration change’ on information processing. For an oscillatory system, when a perturbation occurs at a specific phase of its cyclic activity, the following oscillatory activity is shifted by a consistent amount. This can be quantified by a phase-response-curve (PRC) [42, 39, 13] by tabulating the phase shift Δ*ϕ* induced by a perturbation in dependence on the phase *ϕ* at pulse onset (Fig. 2A). Conversely, a PRC can be used to determine the ‘right’ time for a stimulation in order to shift the system’s phase by a desired amount.

**Figure 2.**
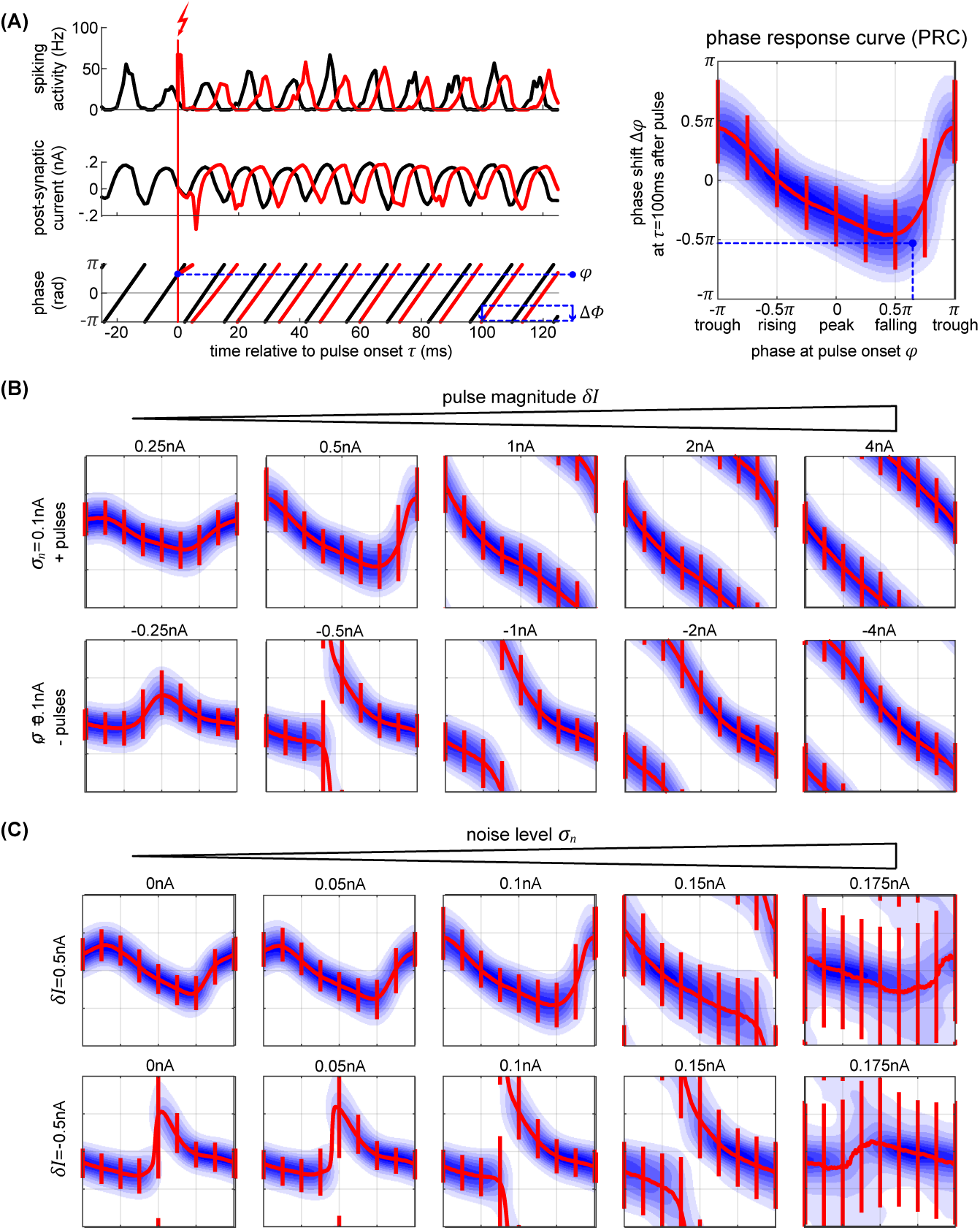
**Phase response curves.** **A.** The plots on the left (top and middle row) show the activity of the excitatory population (in black) and how a single perturbation applied to the oscillator changes both activity and post-synaptic current (in red). From the corresponding phase dynamics (bottom row), we capture the phase *ϕ* at the pulse onset, and the resulting phase shift *δϕ* 100ms later (blue arrow). This gives us a single data point (marked in black) in the PRC space on the right. By repeating this procedure we obtain the distribution of pulse responses indicating the PRC and its variability due to internal and external noise sources. **B,C.** We show multiple PRCs across different conditions - varying pulse magnitude (B) and internal noise level (C). In each plot, the thick red line represents circular mean of the phase shift across the pulse onsets, while the thin red lines indicate the corresponding 25th and 75th percentiles. At low pulse strengths, the resulting PRC shows a smooth biphasic relationship - pulsing at the peak results in a negative phase shift (delay) and pulsing at the trough gives a positive phase shift. As we increase the strength of the perturbation the magnitude of the phase shift increases. At sufficiently high pulse magnitude of either polarity, a perturbation leads to a complete phase reset.

We simulate electric stimulation by injecting a square pulse of current of 1 ms duration into all the neurons within the oscillator. The pulse was intended to emulate intracortical microstimulation (ICMS), affecting the population of local neurons indiscriminately. We tested depolarizing (positive pulses, exciting the neurons) and hyperpolarizing (negative pulses, inhibiting the neurons) pulse polarities at multiple pulse strengths *δI*, from 0.25 nA up to 4 nA. To collect the PRC curve data, first, we run the model for a total time *T* without any stimulation pulses (Fig. 2A, left panel, black curves). Using real-time phase measurement, this data allows us to tabulate normal, unpulsed phase progression *ϕ*(*t*). For assessing the impact of perturbation on phase, we again run the model for time *T*, now pulsing at random points in time, and obtain the pulsed phase progression *ϕ_δI_*(*t*) (Fig. 2A, left panel, red curves). The resulting phase shift observed after a delay time *τ* is then given by Δ*ϕ*(*τ*) = *ϕ_δI_*(*tonset* + *τ*) − *ϕ*(*t*_*onset*_ + *τ*). Repeating the procedure for sufficiently many *t*_*onset*_’s finally yields – since the dynamics is stochastic – a phase-response curve probability density function *ρ_τ_*(Δ*ϕ|ϕ*) (Fig. 2A, right panel, blue shading). By taking the circular mean across Δ*ϕ*, one can condense *ρ_τ_* into a mean PRC 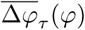 (Fig. 2A, right panel, red line), whose inverse gives the appropriate onset phase(s) *ϕ* which achieve(s) on average a phase shift Δ*ϕ* at time *τ* after giving the pulse.

Note that this inverse mapping does not have to be unique, nor does it have to exist for any desired phase shift, especially for low pulse strengths. In theory, one can realize any desired phase shift by using a sequence of (small) shifts into the right direction, but since we have to cope with a noisy dynamics inducing frequency jitter, we typically aim at achieving a desired shift with as few pulses as possible.

Note that the phase shift Δ*ϕ* does not occur immediately. Rather, following the stimulation, the network takes time to stabilize and settle back into its normal cyclic activity. In the single oscillator model, it takes around 2-3 cycles (around *τ* = 30 ms) for the network to settle into its new stable phase state. Following this time point, the mean phase shift stays consistent, however, the variability goes up, due to the activities’ intrinsic fluctuations in frequency.

For weak perturbations (*δI* = 0.25 − 1.0 nA pulse magnitude), the resulting phase offsets are small, resulting in a smooth biphasic PRC. The negative and positive pulses cause shifts into opposite directions (Fig. 2B, compare top and bottom rows of first plots on the left). However, as the strength of the pulse increases, the phase-shifts increase as well, until they look the same and a complete phase reset occurs resulting in a PRC that approaches the shape of a straight line (Fig. 2B, plots on the right).

As we increase the internal noise of the model, the variability of the PRC goes up with it. At a sufficiently high noise level, *σ*_*n*_ *≥* 0.175 nA, we no longer achieve stable or predictable phase shifts (Fig. 2C), which means that the oscillator has become too chaotic to exhibit phase-response properties [26]. Additionally, when applying a specific magnitude of a pulse, its effect seems to increase (getting closer to a full phase-reset) with noise as well. In part, this is due to the fact that the amplitude of the oscillations in the noisy conditions are lower, meaning that the relative magnitude of the pulse to the oscillations gets higher with increasing noise.

#### 3.1.3 Controlling oscillations

##### Phase control procedure

To test the ability to use the phase-response characteristics of our model, we employ the following task: we run two independent oscillators, X and Y, simultaneously. If we let them run without interfering, the phase difference between their activities Φ_XY_ = *ϕ*_X_ − *ϕ*_Y_ performs a random walk, as their frequencies fluctuate independently from each other^1^. We want to pulse Y to keep it synchronized with X. To achieve this, first, we track their phases *ϕ*_X_ and *ϕ*_Y_ in realtime, using the AR model to forecast signal at each time point (see Fig. 3A). Once Φ_XY_ surpasses the allowed threshold level (more than an eighth of a cycle difference, |Φ_XY_| > *π*/4), we apply a stimulation pulse at just the right phase in order to enact a shift in X’s phase Δ*ϕ*_X_ that is as close as possible to the required correction −ΔΦ_XY_. As soon as *ϕ*_X_ matches the desired onset phase, the stimulation current is given. After the pulse, we enforce a refractory period of *τ*_ref_ = 100 ms when no pulses are allowed in order to let the network settle and maintain its new phase relationship.

**Figure 3.**
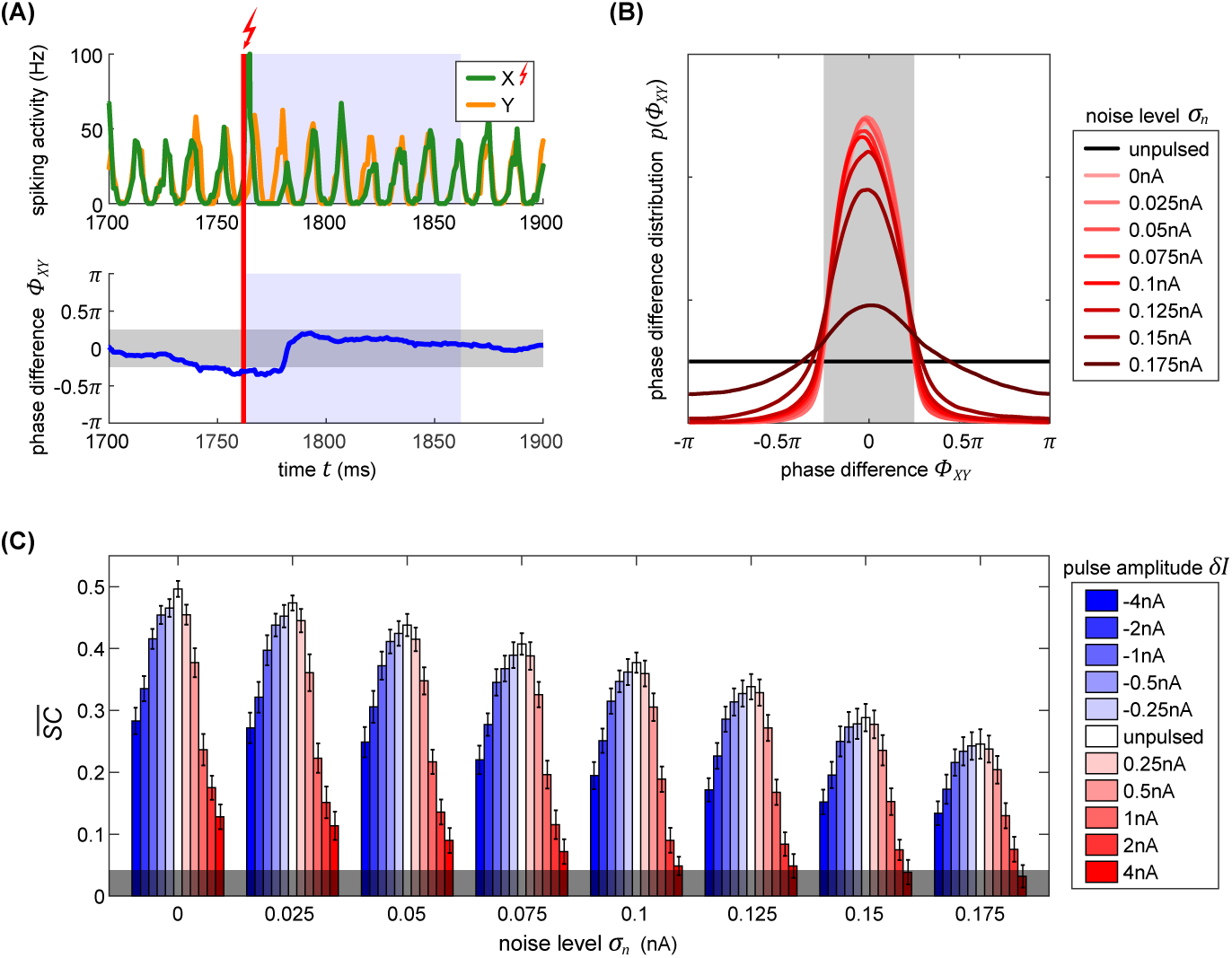
**Using PRCs to synchronize two independent oscillators.** **A.** The top plot shows the activity traces of two independent oscillators, X (orange) and Y (green). As soon as the absolute phase difference between the two oscillators (lower plot, blue line) exceeds the threshold of 0.25*π* (region shaded in gray), the appropriate phase to pulse X in order to achieve the required shift to bring it back into synchronization with Y is determined. Once X is at this ‘right’ phase, a pulse is applied (vertical red line). Following the stimulation pulse, a pulse refractory period is induced for 100ms (shaded blue regions). **B.** Evolution of X-Y phase difference distribution with increasing noise. With higher noise, the amount of time that the model spends in the desired state diminishes, as visible by the broadening distribution. The strength of the applied pulse is much less significant and does not affect the desired state proportion nearly as dramatically (not shown). **C.** Signal content 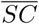 for different pulse strengths (blue-red scale) across varying internal noise conditions (horizontal axis). Stronger pulses cause larger and longer-lasting artifacts in activity which greatly reduces the signal information content measure 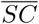. Note that delivering inhibitory pulses consistently leads to activity that contains more stimulus content than excitatory pulses of equal magnitude.

##### Synchronizing two independent oscillators

In Fig. 3B we show the resulting phase difference between X and Y, with the desired state shaded in grey. In the unpulsed control condition, due the inherent variability in the oscillator’s frequency, their phase difference constitutes a random walk, resulting in a uniform distribution. Once the closed-loop procedure is applied, the difference of phases between the oscillators shows the desired distribution centered around the target phase state (-*π/*4 to +*π/*4). Notably, the strength of the pulse does not affect the distribution (hence, just one distribution shown for all pulse strengths in Fig. 3B). The model’s inherent noise level plays a major role. As the noise increases, the ability of the pulsing procedure to maintain the desired state goes down. At the highest noise level, even though there isn’t any sort of a reliable PRC curve, we also achieve a distribution centered around the desired phase difference. Note, that this is not due to any phase-response properties but is merely the effect of applying pulses to the network whenever it is not in the desired state, thus pushing it away from the ‘forbidden’ state, and then letting it run passively whenever the desired state is achieved - the phase onset of the pulse does not matter. This serves as a good control to compare the other curves with.

Next, we use spectral coherence to assess the amount of input signal information that is present in the networks’ output activity (Fig. 3C). Not surprisingly, by pulsing the population, we are degrading the stimulus content. With perturbations of higher magnitude, the amount of degradation increases appropriately. Notably, negative perturbation pulses (in blue) result in significantly less information degradation than the positive pulses (in red). At high noise levels and at a sufficiently high pulse magnitude (4 nA), the amount of stimulus information is no longer significant and falls below the 95% confidence interval at the bottom of the plot. In summary, if we want to use electrical stimulation for assessing information processing in the brain, we have to take care to use an appropriate pulse strength for not completely masking the signals whose representations we are targeting.

### 3.2 Part II: Bistable columnar network

Here the techniques developed in the first part of our study will be applied to an established, prototypical columnar network implementing selective signal routing under attention. After briefly describing the model itself and its dynamics, we will first quantify how the model reacts to perturbation pulses applied to different parts of the system. Using this knowledge, we can finally interact with the model ‘cortex’ in a meaningful way, simulating the effects of ‘natural’, physiological attention by using ‘artificial’ pulses to selectively route external signals to neural target populations. Conversely, our results provide predictions which can be used in physiological experiments to specifically test the particular model setup and, on a more general level, hypotheses about the still debated neural mechanisms realizing communication-through-coherence.

#### 3.2.1 Structure and dynamics of columnar network

##### Setup and connectivity

We use the cortical column setup from Part I to construct a model composed of several interconnected oscillator modules, representing interactions between cortical columns in areas V1 and V4, similar to the work of [27]. All the projections between the cortical columns originate from their respective excitatory subpopulation, reflecting the finding that inhibitory neurons have been found to form primarily local connections, whereas the excitatory neurons project to up- and downstream visual areas [44], and laterally to neighboring columns [45].

A schematic of the model is presented in Fig. 4A. The input (upstream) layer of the model is composed of two oscillators, X and Y, representing two neighboring V1 cortical columns. These are driven by afferent connections delivering independent Poisson spike trains, each modulated by its own input signal, S_X_ and S_Y_. Furthermore, X and Y share a connection from the excitatory pool of neurons of one population to the inhibitory neurons of the other, X_*e*_ to Y_*i*_ and Y_*e*_ to X_*i*_ with connection probability *p*_XY_ = 0.02 and delay *τ*_XY_ = 5 ms.

**Figure 4.**
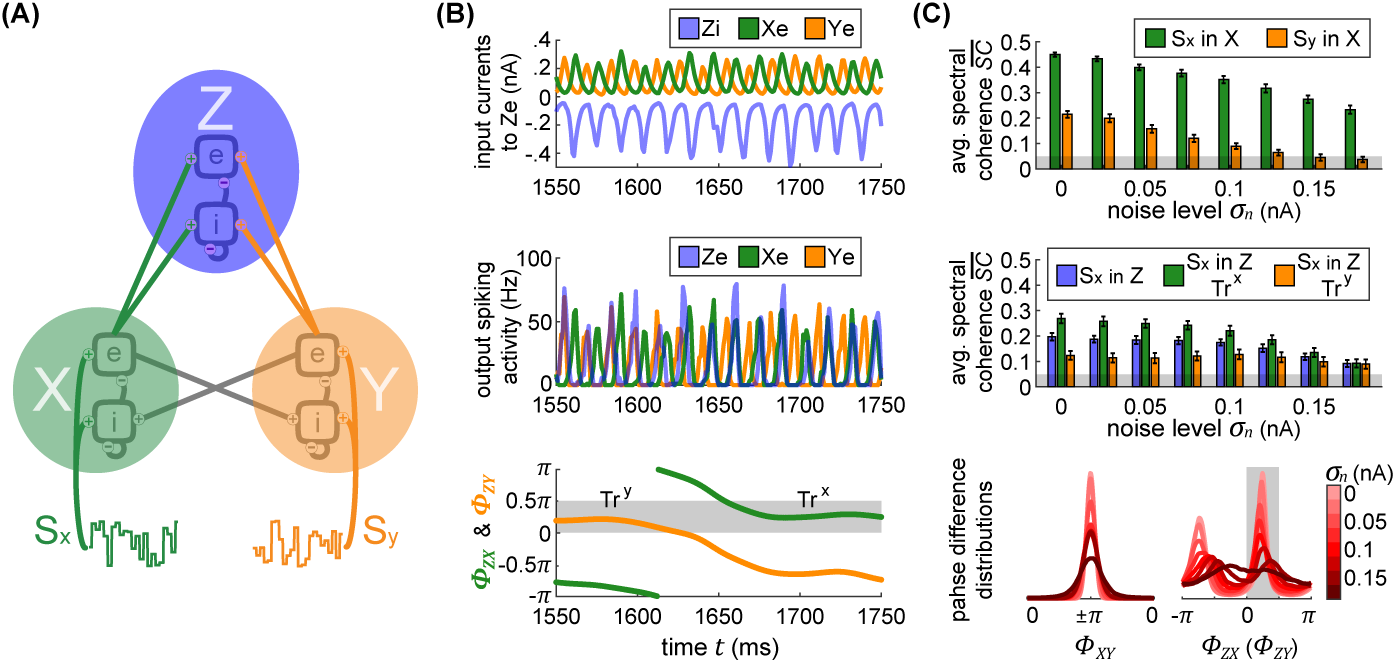
**Bistable XYZ model.** **A.** The columnar network consists of three cortical populations X, Y and Z. X and Yform the lower layer of the model, and each population is driven by its own input signal *S*_*x*_ and *S*_*y*_. X and Y are connected by lateral interactions from the excitatory sub-population of one column to the inhibitory population of the other column. The lateral connectivity is set up such that the two populations establish an antiphase relation. The output of X and Y serves as input into population Z forming the upper layer. **B.** The top plot shows all the postsynaptic currents in Z_*e*_ as a result of the inputs it receives from X_*e*_ (green), Y_*e*_ (orange) and Z_*i*_ (blue). The middle plot shows the spiking outputs of all three populations, in the same colors as above, and corrected for the synaptic delays between the populations. The bottom plot shows the phase differences between Z and X activity Φ_*ZX*_ (orange) and between Z and Y activity Φ_*ZY*_ (green) with a shaded region representing a favorable phase state for either population. Note that X and Y’s activity is consistently in antiphase. **C.** The top plot displays the input signal content 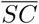 in the activity of X and Y as a function of internal noise level. Due to lateral connectivity between the two populations, there is slight signal mixing, hence X activity manifested mostly the input signal *S*_*x*_ (green) and significantly less of input signal *S*_*y*_ (orange). As the noise grows higher, both signal contents decrease. The middle plot quantifies routing of each input signal through to Z. The blue bars show the content of either one of the input signals in Z regardless of the current state of the network. Conversely, the green and orange bars display signal content of S_X_ when the network is in a Tr^X^ or in a Tr^Y^-state, respectively. The bottom plots show distributions of phase differences for different levels of internal noise (red color scale) for the X-Y populations (left) and X-Z (Y-Z) populations (right). The two peaks of the distributions in the right-hand plot indicate a bistable dynamics of the network. Phase differences between 0 and 0.5*π* correspond to the preferred state when information routing should be optimal (shaded in grey).

In the output (downstream) layer of the model, a third oscillator Z represents a single V4 cortical column that receives input from each of the V1 populations, emulating the convergence of receptive fields when going downstream in the visual system. X_*e*_ and Y_*e*_ project with equal strength onto Z_*e*_ with connection probability *p*_Z*e*_, and onto Z_*i*_ with connection probability *p*_Z*i*_.

For different noise magnitudes *σ*_*n*_, we scale the driving magnitude of the afferent inputs into each population in order to maintain consistent firing rates across the conditions. First, *S*_*e*_(*t*) and *S*_*i*_(*t*) are scaled for the X and Y oscillators, similar to the case with a single cortical column. Once X and Y evoke the desired output spiking rate of 15 Hz, *p*_Z*i*_ and *p*_Z*e*_ are scaled such that Z is sufficiently driven to display the same spiking rates as well.

##### Model dynamics

Each of the individual oscillators retains the parameters of a single cortical column from before, resulting in cyclic activity in the gamma range for each of the populations. In the top row of Fig. 4B, we display all the input currents to the Z_*e*_ population, which receives inputs from X_*e*_, Y_*e*_ and Z_*i*_. Due to the intra-population connectivity between X and Y, their oscillations are consistently in an anti-phase relationship. The inputs received from X and Y compete to entrain Z, which results in bistable model dynamics as described in [27]. In effect, when Z is entrained by X, the troughs of Z_*i*_’s input to Z activity correspond to the peaks of X and to the troughs of Y (and vice versa when Z is entrained by Y). Due to the combined effect of all the noise sources, Z does not get stuck in either stable state, but rather switches stochastically between the two.

The bistable dynamics of the system are clearly visible in the bottom row of Fig. 4C where we plot a histogram of Z-X and Z-Y phase differences, Φ_ZX_ and Φ_ZY_, respectively, across multiple noise conditions. As can be expected, with increasing levels of noise, the preference for either stable state grows weaker.

##### Signal transmission

The inherently bistable dynamics provides a perfect mechanism for implementing communication through coherence (CTC). The CTC hypothesis states that when a population receives multiple oscillatory inputs, it can selectively route one and suppress the others by establishing favorable and unfavorable phase relationships, respectively. For example, in the first half of the trial shown in Fig. 4B, X input to Z_*e*_ arrives when Z_*e*_ is least inhibited by the Z_*i*_ input, putting it into an excitable state and allowing the information content of the signal in X to propagate into (and through) Z. Simultaneously, the bursts of Y’s activity arrive concurrently with maximal inhibition from Z_*i*_, hence suppressing Y’s information content. In sum, the output spikes of Z during this period primarily reflect the activity it receives from X. The same is true in the other direction – when Y wins the entrainment ‘battle’ over Z, its output propagates onwards, while X’s output is effectively suppressed. In the following, we will call these two stable states trans-X-favorable (abbreviated Tr^X^) and trans-Y-favorable (abbreviated Tr^Y^).

By using the spectral coherence, we assess the content of each input signal, S_X_ and S_Y_, in all three populations X, Y and Z. Due to the recurrent connections between X and Y that were not present in the independent case considered in part I, there is a weak mixing of the input signals in the first layer, as seen in the top plot of Fig. 4C. X represents primarily S_X_ (orange line), and to a small but significant extent S_Y_ (green line). For reasons of symmetry, the same lines also represent signal content in Y (orange for S_Y_ in Y, and green for S_X_ in Y).

If we compute the representation of each input signal in Z output without regard for the current state (Tr^X^ or Tr^Y^), we find that on average each signal is equally expressed, as indicated by the blue bars in the middle plot of Fig. 4C. However, when we assess signal content only at times where the network is in the Tr^X^ state, we find that Z activity contains significantly more information from S_X_ (orange bars in Fig. 4C) as opposed to S_Y_ (green bars in Fig. 4C). This result indicates that the model does indeed perform signal routing, while stochastically switches between the two equivalent input sources. As we increase the noise, qualitatively, none of these relationships change – populations in a favorable phase relation always route more information. However, at a sufficiently high noise level (*σ*_*n*_ *≥* 0.15 nA), the difference between S_X_ and S_X_ signal routing through Z is no longer significant.

Note, that the network’s bistability states do not correspond to perfect zero-phase or anti-phase synchronization. In part, this is due to the slight deviations of the neural signals from a perfect sinusoidal oscillation which distorts phase extraction. Further, the dynamics of the model themselves appear to work in a way such that the Z_*i*_ current oscillations occur shortly before the peaks of either X_*e*_ or Y_*e*_ currents. With this in mind, we mark the phase difference from 0 to 0.5*π* as the preferred phase state for signal routing (shaded in grey in Fig. 4B and C).

#### 3.2.2 Pulse-response characteristics of the columnar network

As introduced earlier, we will use *ϕ*_*i*_ to denote the phase of oscillator *i*, and Δ*ϕ*_*i*_ to denote the change in phase of oscillator *i* induced by an external pulse. The probability *ρ_τ_*(Δ*ϕ*_*i*_|*ϕ*_*i*_) to observe a phase shift Δ*ϕ*_*i*_ a delay *τ* after a pulse was given when the oscillator was at phase *ϕ*_*i*_ then describes the (stochastic) *phase response-curve* (PRC) of unit *i*.

However, in our extended model an oscillator is part of a network in which a single oscillator’s phase might be less important than *phase differences* between *pairs* of oscillators. For example, in order to gate an input signal from population X to population Z, their phase difference must be close to 0 as was observed in the previous section. For this reason, we will also consider how the phase difference Φ_*ij*_ = *ϕ*_*i*_ − *ϕ*_*j*_ between populations *i* and *j* is affected by a pulse, giving us a distribution *ρ_τ_*(ΔΦ_*ij*_|Φ_*ij*_) over induced phase difference shifts ΔΦ_*ij*_ which we will term *phase-difference response curve* (PDRC). One can distinguish two conceptually different possibilities to interact with the columnar network: Pulsing population X (or Y) from the input layer, or pulsing population Z in the output layer. In the following paragraphs, we will investigate these two possibilities in more detail, with Fig. 5 illustrating the corresponding effects at a delay of *τ* = 100 ms after the pulse, at an intermediate noise level of *σ*_*n*_ = 0.075 nA.

**Figure 5.**
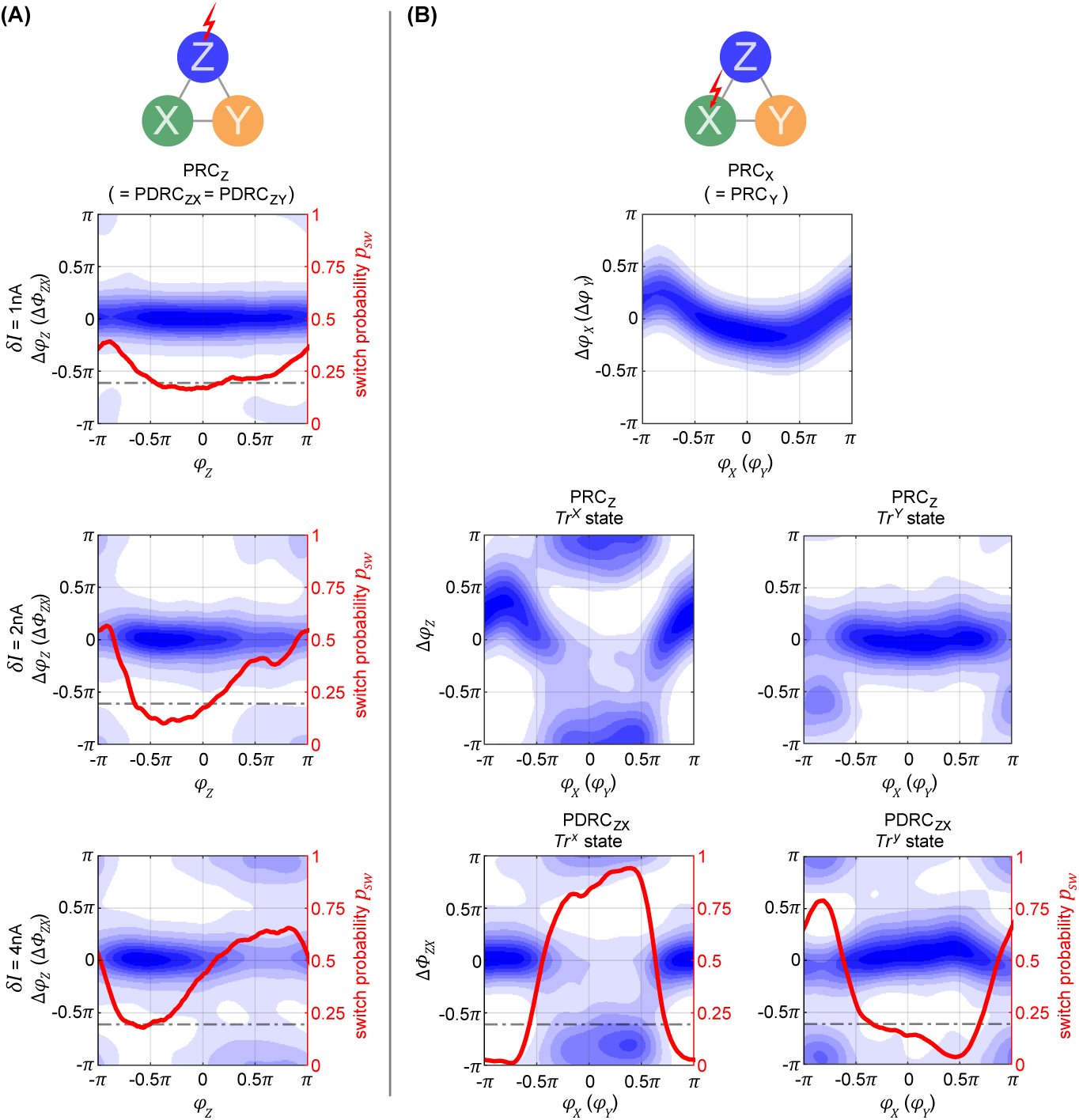
**Phase-response properties of the XYZ model.** **A.** Phase-response curves for pulsing Z with varying magnitudes, sampled at 100ms after pulse onset. Since Z does not send any projections to X or Y, the same plots also represent the *phase-difference* response curves for Φ_*ZX*_ and Φ_*ZY*_. The distributions are centered around zero (phase state does not switch) and *π* (phase state switches). The red lines show the state switch probability *p*_*sw*_ across the different onset phases *ϕ*_*Z*_. **B.** Phase response characteristics when X is pulsed with a current of *δI* = 0.5nA, 100ms after the pulse onset. The top plot shows the PRCs for X and also for Y, since they both end up shifting by approximately the same amount because of maintaining their anti-phase relationship. In the middle row, the two plots show the resulting phase shifts Δ*ϕ*_*Z*_ in Z depending on the phase state *ϕ*_*X*_ at pulse onset. Once we take the phase difference between the corresponding data points in the top plot and the data points in the middle plots, we obtain the phase-difference response curves displayed in the bottom row. Note that the distributions are centered around zero, indicating the network staying in the same state, and around *π*, indicating that the pulse causes the network to switch states.

##### Pulsing output layer population Z

Since Z does not send feedback projections to the input layer, the effect of a pulse stays confined exclusively to Z’s activity and is independent on the system being in state Tr^X^ or Tr^Y^. Because of this, any phase shift Δ*ϕ*_Z_ induced onto Z is equivalent to the shifts in phase difference ΔΦ_ZX_ = ΔΦ_ZY_ = Δ*ϕ*_Z_ between X and Z, and between Z and Y. In Fig. 5A we can see that, independently on noise magnitude, the resulting PRC and PDRC distributions are bimodal and have peaks around 0 and *π*, unlike the effect of a pulse on the single cortical column studied in the previous section. This result is due to the bistable dynamics, which after the immediate effect of the pulse cause Z’s phase to continue shifting until one of the stable states is reached. Accordingly, the final phase shift can be either close to zero or close to *π*, corresponding to no system state change or to a switch between stable states Tr^X^ and Tr^Y^, respectively.

When the pulse magnitude is sufficiently small, e.g. *δI ≤* 1 nA, a state change is unlikely (Fig. 5A, top plot) since the corresponding average phase shift for a single oscillator is too small, Δ*ϕ*_X_ *≤* 0.5*π*. Once we increase the pulse strength (Fig. 5A, middle plot), the likelihood for a phase shift of *π* increases, with their respective phase onset locations roughly corresponding to the ones which led to Δ*ϕ ≥* 0.5*π* phase shifts in the single oscillator (e.g. compare to Fig. 2B, bottom middle plot). In order to best summarize the concentrations in the PDRC densities around 0 and *π*, we calculate the state switch probability *p*_sw_ across the onset phases *φ*_z_ via 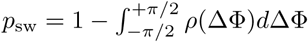. Increasing the pulse strength beyond 2 nA affects which phase onsets correspond to a higher switch probability, but does not seem to affect the minimum and maximum switch probabilities (Fig. 5A, middle and bottom plots).

##### Pulsing input layer population X

If we apply the pulse to one of the lower level populations, the perturbation will propagate forth and back via recurrent connections and lead to a cascading effect of pulse echos. However, after more time passes (*τ* = 100 ms is more than sufficient), the X–Y populations settle back into their anti-phase relationship. Because of this, at a sufficient delay *τ*, the PRC densities for X and Y are essentially identical and appear to resemble a diffused version of the PRC in the single oscillator case (compare the top plot from Fig. 5B to the corresponding plot in the middle of Fig. 2B).

For understanding how a pulse delivered to X affects the phase of population Z, we have to consider whether the system is in state Tr^X^ or Tr^Y^. When in Tr^X^, the propagated pulse arrives at Z at about the same phase as the initial perturbation was given to X, resulting in the PRC density shown in the left plot in the second row of Fig. 2B. When in Tr^Y^, a pulse given at a peak of X’s activity now arrives at a trough of Z’s activity, giving rise to a different PRC as shown in the right plot of the second row of Fig. 5B.

How do these different effects of a pulse given to X combine and affect the global state of the columnar network? To obtain the corresponding measure ΔΦ_ZX_, we can take the difference between the corresponding data points from the PRC of X (Fig. 5B, first row) and the PRC of Z (Fig. 5B, second row). The resulting PDRC densities are displayed in the bottom row of Fig. 5B, revealing again a bimodal distribution with peaks at 0 and *π*. Notably, even though the initial pulse magnitude of 0.5 nA was insufficient to obtain a high state switching probability when pulsing Z, when pulsing X with the same strength, for a large range of pulse onset phases a much higher switching probability is observed (red curves in bottom row of Fig. 5B). There are two reasons why switching becomes easier: first, the perturbation does not only affect one population but is propagated to all other ‘players’ in the network, and second, for a brief period of time after the pulse, anti-phase relationship between X and Y is affected.

#### 3.2.3 Optimizing stimulation pulses for state switching

Our goal of using stimulation is to cause the network to be continuously in a desired state Tr^X^ or Tr^Y^ for either transmitting signal X or signal Y, respectively. By deriving state switching probabilities from phase-response curves as described in the previous subsection, we now have a tool for optimizing the stimulation pulse parameters towards this goal. Accounting for symmetry between X and Y, all the following results are presented with the aim of switching to a Tr^X^-favorable state. By this design, whenever the network is already in a favorable Tr^X^ relationship, no perturbation is necessary. However, if at any point the network instead is in a Tr^Y^ favorable relationship, we can apply a pulse either to column X, Y, or Z to attempt to switch the state to Tr^X^.

In Fig. 6, we display the columnar network’s state switch capabilities for Tr^Y^ *→* Tr^X^ for negative (left major column) and positive (right major column) pulses, and for the three possible target columns X, Y, and Z of the pulse (upper, middle, and bottom rows, respectively). The panels for each of these six conditions are organized in groups of three by three graphs, displaying the switching characteristics across multiple noise levels (columns) and multiple pulse magnitudes (rows). In particular, since the unpulsed system possesses a non-zero passive switch probability, we plot the pulse-induced *change* in switch probability in dependence of the onset phase *ϕ* of the pulse (horizontal axis), and in dependence of the time *τ* following the initial pulse (vertical axis). Thus, negative values (blue shading in the plots) indicate that, if delivered at the wrong moment, perturbations can actively *hinder* a transition to Tr^X^ and instead stabilize the undesired Tr^Y^-state. In the following, we will briefly discuss the effects of pulsing the different target columns.

**Figure 6.**
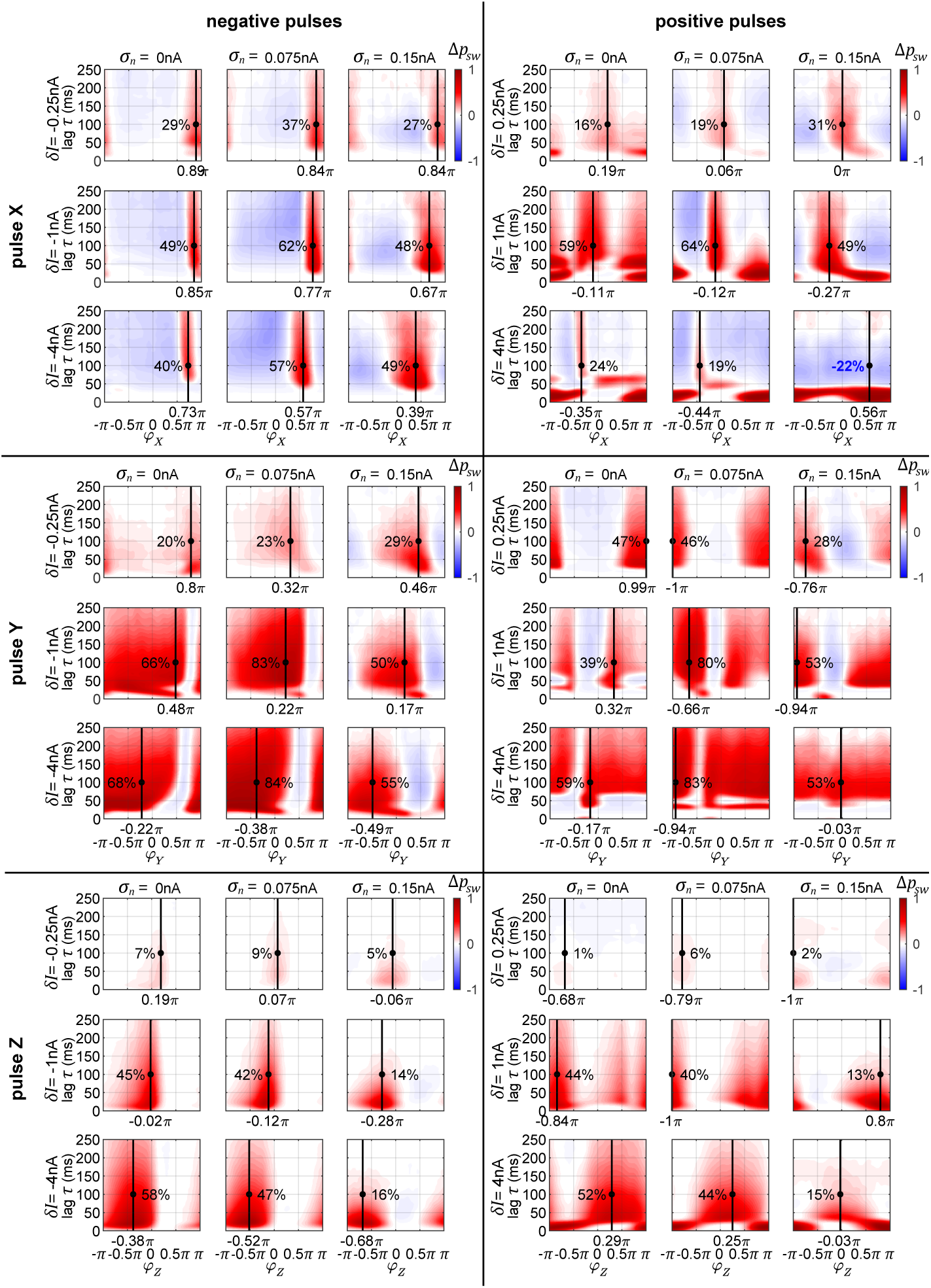
**Changes in state switching probability.** State switch capabilities for a transition Tr^Y^ *→* Tr^X^ for negative (left major column) and positive (right major column) pulses, and for the three possible target columns X, Y, and Z of the pulse (upper, middle, and bottom rows, respectively). The panels for each of these six conditions are organized in groups of three by three graphs, displaying the switching characteristics across multiple noise levels (columns) and multiple pulse magnitudes (rows). The color codes for the pulse-induced *change* in switch probability in dependence of the onset phase *ϕ* of the pulse (horizontal axis), and in dependence of the time *τ* following the initial pulse (vertical axis).

##### Pulse X

The graphs in Fig. 6, top row, reveal that in addition to having an optimal onset-phase, for each noise condition, there is also an optimal pulse magnitude that results in the largest increase in switching probability. Interestingly, a medium level of noise results in larger switch probabilities than the zero-noise condition.

When applying a negative pulse, there are always intervals of phase onsets that increase, and intervals that decrease the probability of the network switching its state. On the contrary, when applying a positive pulse, at a high noise level (*σ*_*n*_ = 0.15 nA) and a high pulse magnitude (*δI* = 4 nA) the onset phase does not appear to matter for the final outcome. In this particular case, all phase onsets lead to a decrease in the switch probability. If we would utilize an even stronger pulse, we would also see this effect when the network noise level is low.

The amount of time it takes the network to settle down onto a new state tends to increase with pulse strength (*≈* 30 ms for *δI* = 0.5 nA pulse vs. *≈* 60 ms at *δI* = 2 nA and 4 nA). A stronger initial current causes a stronger reverberation of the perturbation, which then takes longer to decay within the system, increasing the time it takes the columnar network to settle back to its normal activity. Once a system converges onto a stable state, the switching probability change decreases due to the passive stochastic switch chance (going up on the vertical axis). This effect is particularly visible for the positive pulses, where we get to observe the different phase-states the system goes through before settling down.

##### Pulse Y

When pulsing Y instead of X, the graphs in Fig. 6, middle row, reveal that the switch probabilities appear complementary to the ones from pulsing X. When pulsing X, if a specific phase onset leads to an increase in switching probability, the same phase onset typically leads to a decrease in switching probability if pulsing Y instead. This makes sense, since by changing which population (X or Y) we are pulsing at one specific pulse onset phase, we are essentially changing the onset phase of the propagated pulse that arrives to Z by an amount of *π*, since X and Y maintain an anti-phase relationship.

Because of this, when applying a positive pulse of a large magnitude (*δI* = 4 nA) in the noisy condition (*σ*_*n*_ = 0.15 nA), the probability of a switch is now consistently high across all pulse onsets, whereas in the previous condition a pulse to X was always decreasing switch probability.

##### Pulse Z

As described previously for Fig. 5, when pulsing Z, a pulse of low magnitude is hardly sufficient for inducing a significant change in the switch probability. As the noise level of the system increases, the switch probability decreases substantially (Fig. 6, bottom row).

#### 3.2.4 Using stimulation pulses to control signal transfer in the columnar network

The paradigm to control the synchronization state of the network is similar to controlling the phase of an independent oscillator, with one crucial difference: In the independent oscillator case, a pulse is applied at various phase onsets, depending on what sort of a phase shift is currently necessary. In the columnar network, however, the choice is binary: to switch or not to switch. If we desire to change the current system state, then there is just one specific optimal onset-phase for the pulse. So, for every pulsing condition (i.e., which population pulsed, pulse magnitude, pulse polarity, and network noise level), the state control procedure comes down to the following:

At a time point *t*, apply the stimulation pulse if the following conditions are met

1. Last pulse was more than *τ*_ref_ ago.
2. The system is in the wrong state and a switch is necessary.
3. The current phase of the pulsed population corresponds to the one that leads to the highest switch probability.

In order to evaluate how well the pulsing procedure works, we first quantify the proportion of time that the system spends in the desired target state (Fig. 7). At the top of each plot, we display the proportion of time that the network spends in that state without pulsing. As defined previously, the desired state is set to the interval Φ_ZX_ ∈ [0, 0.5π], which corresponds to a quarter of the full interval of possible differences.

**Figure 7.**
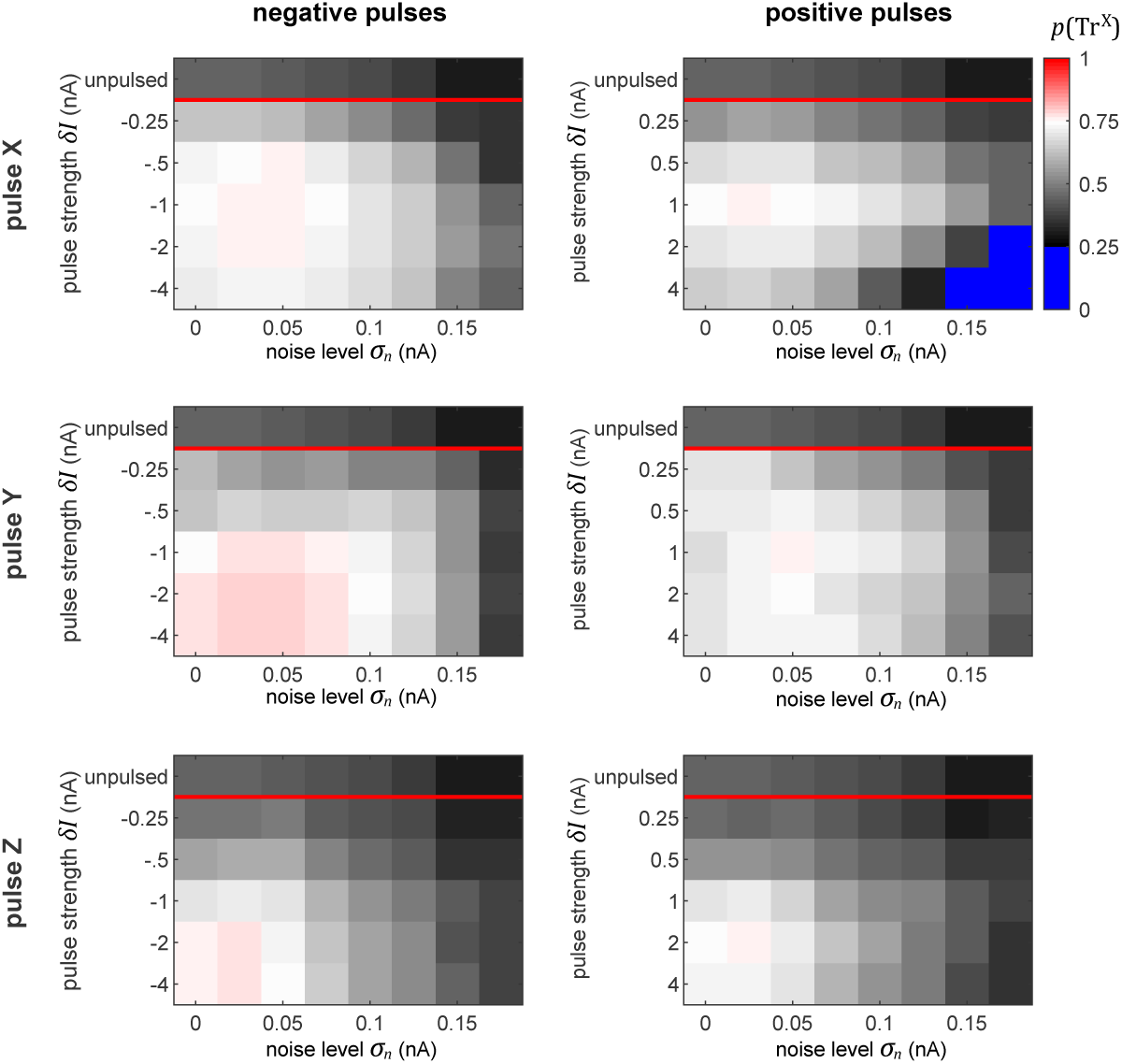
**Proportion of time spent in desired state.** The plots display the proportion of time that the network spends in the desired state for all the conditions, as labeled. The top row in each subplot indicates the amount of time that the network spends in the desired phase state passively, without any perturbation pulses. Note that the color map turns sharply blue below values of 0.25, which corresponds to the chance level of being in the desired state for a uniform distribution of phase differences between sending and receiving populations.

The goal of the perturbation pulses is to increase this value as much as possible. The results observed in this figure perfectly reflect the corresponding switch chance as predicted by the plots in Fig. 6. This is especially clear in the case when we pulse X with using a large positive perturbation (*δI ≈* 4 nA) at a high noise level (*σ*_*n*_ *≈* 0.15 nA). In this condition, the effect of the pulse can only decrease the switch chance, indicated by the blue regions in the top right plot. In all other cases, the procedure succeeds at increasing the amount of time the network spends in the desired state.

Similarly to our previous results from pulsing independent oscillators, the performance of the procedure decreases with increasing noise level. On the contrary, in the columnar network the pulse magnitude and polarity plays a crucial role, whereas in the independent case, the strength of the pulse had no significant effect on the performance. In fact, we observe qualitatively different patterns for which pulse is optimal across the different pulse-polarity and pulsed-target conditions. For instance, when pulsing X, a pulse of *δI* = −1 nA or 1 nA achieves the best performances. However, when pulsing Y, negative pulses get better results at higher magnitudes (saturating at sufficiently high levels), whereas positive pulses exhibit lower performance once the magnitudes are sufficiently high.

Generally, with our model’s specific setup, the results seem to indicate that pulsing Y (i.e. the population whose information we wish to suppress) provides a much more robust and forgiving conditions, by having more admissible phases of the perturbation onset that result to a state switch.

Further, we evaluate the signal routing performance of the pulse procedure by evaluating the difference between S_X_ and S_Y_ signal contents in Z. These differences are displayed in Figure 8. At the top of each plot, the maximally achievable difference is displayed, as seen in Fig. 4C, by evaluating the signal content in Z without delivering any pulses, but for the time intervals in which the system is spontaneously in the preferred state for transmitting a specific signal.

**Figure 8.**
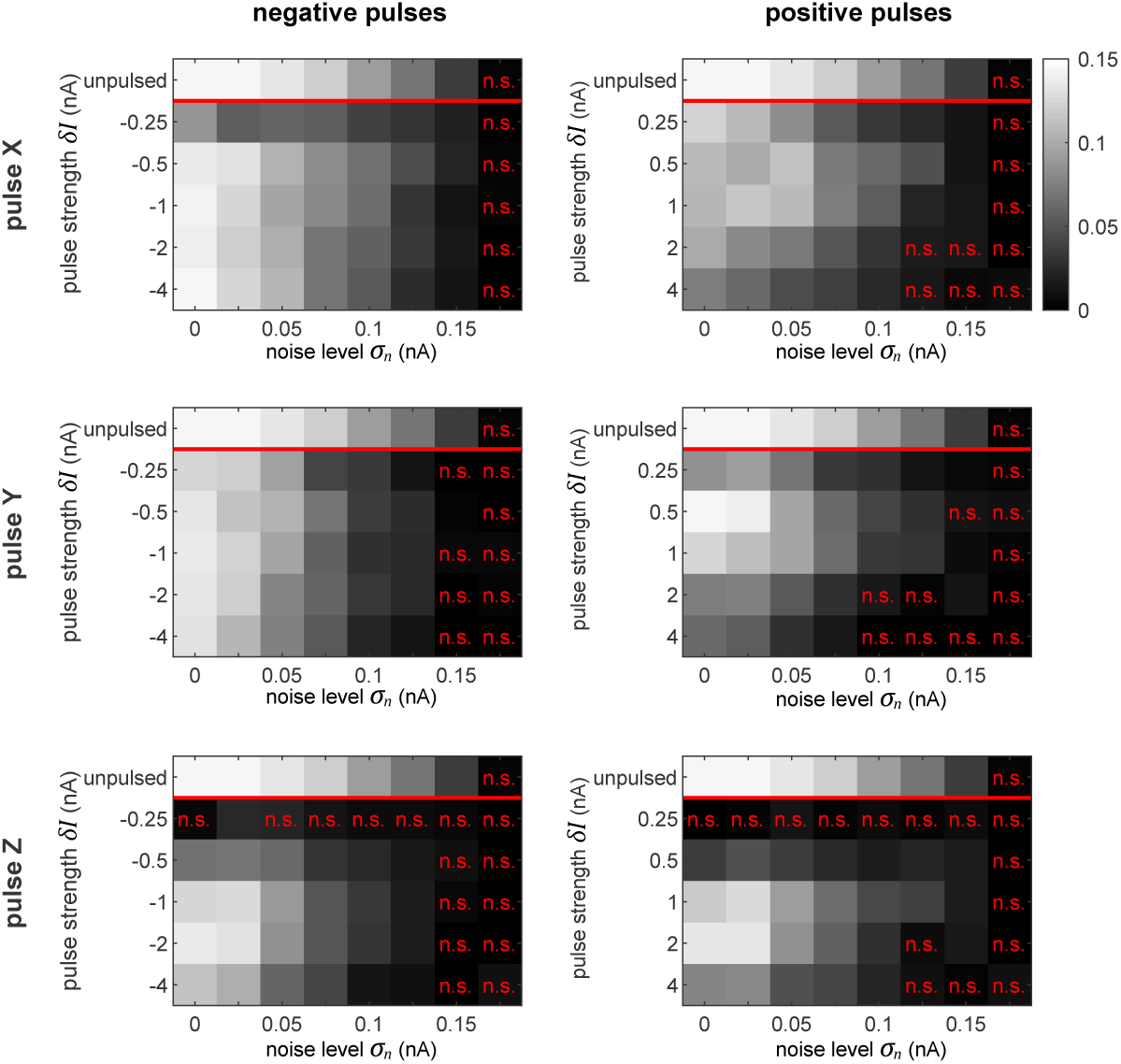
**Selective signal routing via precisely pulsing the columnar network.** Difference in transferred signal content between the two competing input stimuli S_X_ and S_Y_ for different stimulation paradigms (rows) and pulse polarities (columns), evaluated for different pulse magnitudes *δI* (vertical axes) and noise levels *σ*_*n*_ (horizontal axes). The top row in each graph shows the corresponding ‘optimal’ or maximally achievable signal routing performance extracted from time intervals when the network was in a particular preferred state (here, in Tr^X^). Parameter combinations marked by a red ‘n.s.’ indicate conditions where the difference in signal transfer was not significantly different from zero (student T-test, *p* < 0.05).

As we observed in the independent oscillator case, increasing the pulse magnitude increases the amount of degradation of the external signal in the population’s activity in all the conditions. In some cases, a pulse of a higher magnitude actually leads to a better performance in terms of keeping the network in the desired state, but simultaneously, it also increases the amount of signal degradation. Since we focus on signal routing differences, the results in Fig. 8 reveal the appropriate compromise to achieve the best gating performance.

In a recurrent network, the effect of the pulse on external signal representations is not straightforward and hard to predict. First, the signal represented by the pulsed population is degraded by the injected current. Subsequently, the pulse propagates throughout the rest of the system, causing further degradation of signal representations in *other* populations. Consequently, the signal routing performance crucially depends on which population is pulsed. For instance, if we compare the results in the zero noise condition when pulsing X or Y with a positive pulse of *δI* = 1 nA, we find that pulsing Y provides much better routing results, even though pulsing X is actually better at establishing the desired network state (see Fig. 7). In this case, a significant part of the result is caused by signal S_Y_ getting degraded substantially more than signal S_X_.

## 4 Discussion

### 4.1 Summary

The goal of this study was to investigate how precise perturbations can control a recurrently coupled neural network by using its natural tendency to be in one of several preferred network states. For this purpose we developed a closed-loop paradigm to monitor the system state in realtime and utilized the results to deliver rare, but accurately timed stimulation pulses of proper magnitude. First, we evaluated the method on a structurally simple system – the model of a single cortical column. Here, it was possible to synchronize two independent oscillators up to a critical noise level, and to determine the optimal pulse strength. Next, we applied our paradigm to a more elaborate and biophysically more realistic network comprising multiple recurrently coupled columns, proposed [27] as a prototypical implementation for selective information processing via communication-through-coherence (CTC) in the visual cortex. For successfully interacting with such a system, our results demonstrate that understanding the behavior of one of its constituents in isolation (e.g. by obtaining the phase-response curves, PRCs) is not sufficient – instead one has to probe the network as a whole, which required to compute phase-*difference*-response curves (PDRCs). Furthermore, we investigated several ways of interacting with the system, targeting either upstream or downstream neural populations. Ultimately, we could simulate the effect of physiological attention and gate signals by bringing the desired population(s) into a preferred (or non-preferred) phase relationship.

### 4.2 Realism of model and significance of results

Certainly the columnar network is still an abstraction of the real networks performing selective information processing in the visual cortex. We only considered three coupled columns, back-projections from downstream visual areas were not modeled, and we assumed a lateral recurrent coupling structure which is still subject to on-going physiological and anatomical research. Additionally, the effects of our perturbation pulses on neural processes are highly simplified in the simulations. However, even when taking these restrictions into account, we believe our work contributes in three important aspects to the field:

- For being successful in interacting with a neural system, the current state of the system *does matter*. This is particularly obvious when trying to construct a visual intracortical prosthesis [31]. Since there is an on-going dynamics in the cortex even in the absence of an actual visual stimulus [3], it is important to know when an artificial stimulus would be most effective, either in inducing a certain percept or in pushing the system into or towards a desired network state. Another requirement is to ensure an ongoing stimulus processing in downstream visual areas. For this purpose, it would be necessary to first bring the network into a state where incoming information can be successfully gated across different stages. This goal was successfully reached in model simulations of our closed-loop stimulation paradigm.
- With respect to selective information processing, we investigate *one specific* of potentially many implementations of the CTC principle. Our results therefore constitute a prediction of how the real network would behave if it would work according to our hypothesis. In particular, we predict that pulsing the column representing the unattended stimulus would be very effective in switching between the different network states and in selectively gating a stimulus. This should not be the case if the recurrent interactions would not push the upstream populations X and Y into an antiphase relation, thus providing an opportunity to test this critical assumption.
- Finally, our study brings together the tools needed to establish realtime control of stochastic neural systems. One important insight for us were the severe restrictions imposed by noise, be it intrinsic or on the observation level. Crucially, we present a paradigm that relies on single and rare stimulation pulses, allowing to network to spend the majority of the time unperturbed. This is in opposition to utilizing continuous stimulation or repetitive pulses that explicitly entrain the system, which we argue results in non-natural and forced brain activity. Thus, we conclude that instead of explicitly forcing a network state, it should be of great benefit to account for the system’s inherent multistable attractor states and utilize the minimal perturbation to let the network settle naturally in a desired network state.

Below, we discuss the relation of the columnar model to experimental data and possible consequences of changes to the model structure for our stimulation paradigm.

#### 4.2.1 Intra-population connectivity

As briefly mentioned in the preceding paragraph, there are two major assumptions in the connectivity between the X, Y and Z modules in the columnar network. First, the connectivity between the lower layer populations X and Y forces them to establish a stable and symmetrical anti-phase relationship between their activities, without establishing a clear winner between the two. Second, there are no back-projections from Z back onto X or Y.

By increasing the connection strength and changing the delays in the X–Y connection it is possible for the populations to synchronize at a different phase, exhibiting the phenomenon of biased competition [34] already in the first layer, and controlling the overall bistability of the model. In such a system, the bistable dynamics are evoked as the two populations switch between which one leads and which one follows. The phase response characteristics of such a scenario of two interconnected oscillators have been thoroughly explored in [56]. If we employed this sort of connectivity between X and Y in our model, the winner of the biased competition in the first layer would also entrain Z. As [56] show, in this scenario one is also able to control the stable state via a precisely timed stimulation pulse.

#### 4.2.2 Local circuitry

The source of gamma frequency oscillations in the brain has been attributed primarily to two mechanisms: ING - interneuron gamma - which we utilize in our model, and PING - pyramidal interneuron gamma [47]. In the ING mechanism, a population of mutually connected inhibitory neurons generate synchronous IPSPs, creating an ongoing rhythm which is then imposed onto the excitatory neurons [55]. In the PING mechanism a volley of excitation stimulates delayed feedback inhibition, resulting in consistent cyclic behavior when the ratio between excitation and inhibition is appropriate. Research shows that both mechanisms can work together to generate gamma frequency oscillations [11, 23, 12, 5]. Either mechanism or a combination of the two constitutes a self-sustaining oscillator and exhibits phase-response characteristics. Hence, we speculate that regardless of the oscillation generating mechanism, the method established in this study can be used to establish desired phase-locking between populations of neurons and route information – however, with a potentially different phase-(difference)-response characteristics.

#### 4.2.3 Transient synchrony

Even at high noise levels, the rhythmic behavior of our system is an idealized version of what is observed in the visual cortex where the amplitude of oscillations, along with the strength of synchronization phenomena occur as transient events that rarely last longer than 100ms. In particular, for the V1-V4 interaction explored in this study, gamma activity tends to occur in bursts at theta frequency through phase-amplitude coupling, corresponding to the rate of attentional sampling [14, 29, 43]. If theta phase amplitude coupling was included in our model, we presume that it should still be possible to control information routing by injecting the appropriate perturbation towards *the beginning* of each theta-coupled gamma burst.

#### 4.2.4 Modeling the perturbation

In the present study, the applied perturbations involved injecting the same amount of current into all the neurons within a local population. This was designed to model the effect of intracortical microstimulation (ICMS). If we wanted to get closer to the true postsynaptic effect of an ICMS pulse, it would be necessary to work out advanced kernels to convolve with the square wave function that we used. Additionally, it would benefit to have different weights of the perturbations effect by neuron type, physical orientation, and distance to the electrode. As long as the final perturbation is sufficiently short and precise relative to the oscillation cycle, the network dynamics should still exhibit a PRC. We believe that the method employed provides a generic pulse that can be easily modified for other potential stimulation techniques. For example, in the case of modeling an optogenetics pulse [56], the stimulation affects just a specific subset of neurons within local population (of type affected by the viral injection of a particular light-sensitive protein).

### 4.3 Outlook

Intracortical microstimulation and other methods of providing ‘artificial’ input to the brain (i.e., optogenetics) are a useful tool for investigating neural information processing in a causal manner. More importantly, these techniques can be employed in brain prostheses, helping patients to compensate for disabilities in vision, hearing and touch. In the extreme periphery, devices such as a cochlea or retinal implant have already been successfully deployed. But what about the next stages in the brain? For example, for patients with a damaged optical nerve, an implant must interface primary visual cortex directly. Here, one would have to cope with on-going processes, feedback from higher areas, and a strong recurrent coupling – the state of the system. Overriding these processes and directly providing the stimulus in a ‘1:1-mapping’ is difficult and could exert substantial stress to the tissue, potentially making long-term applications unfeasible [28]. We propose that one should rather try to swim with the tide, using the natural tendencies of the network as far as this is possible. In the present study, we started to think about the appropriate strategies and methods and tested them on a very simplistic model, designed after the visual system. The logical next steps could proceed into two directions: first, to put these methods to the test by performing animal experiments, and second, to advance on a theoretical and conceptual level by extending the paradigms into space and time, by delivering complex spatio-temporal stimulation patterns appropriate for a system which exhibits a complex spatio-temporal dynamics even in the absence of an actual stimulus.

## 5 Methods

### 5.1 Neurons and synapses

Interactions between neurons are governed by synaptic weights *ω*_*e*_ and *ω*_*i*_ and conductances *g*_*e*_ and *g*_*i*_, which reflect the magnitude and decay speed of EPSCs (excitatory postsynaptic currents) and IPSCs (negative postsynaptic currents) when receiving a spike from an excitatory or inhibitory cell, respectively.

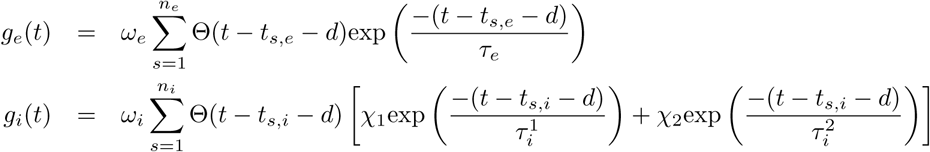

Here Θ is the Heaviside function, *d* the synaptic delay, and *t*_*s,e*_ and *t*_*s,i*_ are the times of presynaptic excitatory and inhibitory spikes, respectively. The decay constants for EPSP and IPSP are given by *τ*_*e*_ and 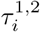, with the inhibitory response containing a mixture of slow and fast components with the relative contributions controlled by *χ*_1,2_. These parameters are set to emulate realistic neurons, in accordance with [4] (see table 1). The activity of the units is simulated in Matlab in discrete time using the forward Euler method with a timestep of *dt* = 0.1*ms*.

**Table 1.** **Neuron and synaptic connection parameters.** The parameters are taken to emulate biophysically-realistic neurons, in accordance with [4]. *p*_0,1,2_ were found by a mathematical reduction of the Hodgkin-Huxley model [1].

|  |  |  |  |  |
| --- | --- | --- | --- | --- |
| **Variable** | $A_e$ | $A_i$ | $p_0$ | $p_1$ |
| value | $2.88e-4 \text{ cm}^2$ | $1.2e-4 \text{ cm}^2$ | $3.89e-9 \text{ A}$ | $1.30e-7 \text{ A/V}$ |
| **Variable** | $p_2$ | $V_e$ | $V_i$ | $V_{thresh}$ |
| value | $1.08e-6 \text{ A/V}^2$ | $0 \text{ mV}$ | $-75 \text{ mV}$ | $-56.23 \text{ mV}$ |
| **Variable** | $V_{reset}$ | $\tau_e$ | $\tau_i^1$ | $\tau_i^2$ |
| value | $-67 \text{ mV}$ | $3 \text{ ms}$ | $1.2 \text{ ms}$ | $8 \text{ ms}$ |
| **Variable** | $\chi_1$ | $\chi_2$ | $\omega_e$ | $\omega_i$ |
| value | $0.9$ | $0.1$ | $0.4 \text{ nA}$ | $1.2 \text{ nA}$ |

### 5.2 Offline phase measurement

For measuring the phase of a cyclic signal, we utilize a well established procedure. For efficiency, the signal is downsampled to 1 kHz and normalized. A power spectrum of the signal is calculated using a Morlet wavelet transform, from which we determine the location and halfway points of the gamma peak. Then, we apply a zero-phase (’filtfilt’ command in Matlab) finite-impulse response (FIR) bandpass filter with bandstops at the halfway points found in the power spectrum. This gives us the gamma component of the signal without distorting the phase. Afterwards, the signal is passed through a Hilbert transform [9], providing us with the complex analytical signal. The argument of the analytical signal gives us the instantaneous gamma phase of the signal. The narrow range of the bandpass filter is necessary, since the instantaneous phase only becomes accurate and meaningful if the filter bandwidth is sufficiently narrow [35].

This sort of a procedure is prone to edge effects and especially to the artifacts induced by sudden spikes in activity due to stimulation pulses used in this study. To decrease the effect of these artifacts, the affected region is set to zero after normalization. Empirical tests showed that the edge and artifact effects on instantaneous phase measurement becomes insignificant around 2 cycles away from the affected region, around 30ms for gamma oscillations.

### 5.3 Realtime phase measurement

When forecasting a discrete time signal *X*_*t*_, given its past time points *X*_*t−i*_, an autoregressive model of order *p* is be defined as

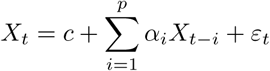

where *α*_1_„*α*_*p*_ are the parameters of the model, *c* is a constant, and *ε*_*t*_ is white noise. The parameters and the magnitude of the noise are trained on a pre-existing set of data using the Burg lattice method (’arburg’ command in Matlab). The model order *p* selection depends on the sampling rate and the characteristics of the input signal and is determined empirically to provide the most accurate phase measurements when compared to the offline phase measurements [33].

For speed and efficiency, the AR model was applied to downsampled 1kHz signals, for which the optimal order *p* was found to correspond to the the average number of time steps within a single cycle of oscillatory activity (approx 15 ms for gamma oscillations). For every condition, a separate AR model is trained on an existing 10 second trial, which provides sufficient amount of data to converge on the appropriate AR parameters.

### 5.4 Information measure significance levels

Chance levels for 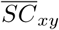 are calculated by pairing up the network activity with 100 surrogate input signals. The resulting distribution of SC values allows us to extract the 95th percentile 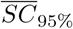, allowing us to evaluate the significance of the information measure score. Further, spectral coherence is affected by sampling size bias. Thus, in order to compare signal routing scores across conditions, they were always computed from 100 seconds of data.

### 5.5 Conditioning of input drive on internal noise

In order to obtain comparable model activity between the different internal noise magnitude model conditions, the input drives, *S*_*e*_(*t*) and *S*_*i*_(*t*) were adjusted to achieve a 15Hz average firing rate for the excitatory and 60Hz for the inhibitory pools of neurons. This was achieved via a simple gradient descent procedure. For example, for a simple oscillator and for the first layer populations X and Y of the bistable model, if the initial drive to the inhibitory neurons led to a firing rate higher (lower) than the desired 60Hz, the inhibitory drive *S*_*i*_(*t*) was decreased (increased) by an amount proportional to the mean-squared error of the firing rate. Simultaneously, if the excitatory pool’s average firing rate was lower (higher) than desired, the excitatory drive *S*_*e*_(*t*) was increased (decreased). The model was then simulated with the updated driving rates and new firing rates were acquired, new firing rate errors were computed and the gradient procedure was repeated until convergence onto the desired firing rate values. In the bistable multi-column model, once the desired firing rates were attained for the X and Y populations, the same procedure was applied to adjust the connection probabilities from X_*e*_ and Y_*e*_ onto Z_*e*_ and Z_*i*_, *p*_Z*e*_ and *p*_Z*i*_.

### 5.6 PRC and PDRC collection details

To collect the PRC curve data, we simulate 2500 runs of one second duration across all the possible conditions (noise level, pulse strength, which population pulsed). In each run, the perturbation occurs at 0.5 seconds, providing us with enough signal before and after the pulse to extract the relevant phase information. In addition, we add a control group where the pulse magnitude is set to 0 - no pulse. For each run, we compute the offline phase across the entire trial. In addition, we determine the instantaneous phase at stimulation onset *ϕ* by using the AR signal prediction procedure to avoid any artifacts caused by the pulse. By pairing up the appropriate trials between the pulsed and the control groups, we calculate the phase difference ΔΦ between the unpulsed and pulsed runs at time *τ* after *tonset*. In the independent oscillator case, the pairing process is only concerned with putting trials together with a minimal difference between their corresponding values of *ϕ* at pulse onset. In the bistable columnar network, the AR procedure is used to evaluate both the onset phase of the pulsed population as well as the phase-state difference Z-X and Z-Y at pulse onset, with the pairing procedure accounting for both, the network state and the stimulation pulse onset phase.

## Conflict of Interest Statement

The authors declare that the research was conducted in the absence of any commercial or financial relationships that could be construed as a potential conflict of interest.

## Author Contributions

DL contributed to the concept of the work, modeling and analysis of data, and draft of the manuscript. UE contributed to the concept of the work, supervision, and draft and revision of the manuscript.

## Funding

This project was supported by the the BMBF (Bernstein Award Udo Ernst, Grant no. 01GQ1106) and by the DFG priority program SPP 1665 (ER 324/3-1).

## Acknowledgments

We would like to thank Daniel Harnack for fruitful discussions in the initial stage of this project and for sharing parts of his program code for simulating the cortical network.

## Footnotes

1 In our mathematical notation, we consistently designate *phases* with *ϕ*, *phases shifts* (induced by the perturbation) with a prefix Δ, and *phase differences* (between oscillators *i* and *j*) with a greek uppercase Φ_*ij*_.

